# Evolution and rapid spread of a reassortant A(H3N2) virus that predominated the 2017-2018 influenza season

**DOI:** 10.1101/543322

**Authors:** Barney I. Potter, Rebecca Garten, James Hadfield, John Huddleston, John Barnes, Thomas Rowe, Lizheng Guo, Xiyan Xu, Richard A. Neher, Trevor Bedford, David Wentworth

## Abstract

The 2017-2018 North American influenza season caused more hospitalizations and deaths than any year since the 2009 H1N1 pandemic. The majority of recorded influenza infections were caused by A(H3N2) viruses, with most of the virus’s North American diversity falling into the A2 clade. Within A2, we observe a subclade which we call A2/re that rose to comprise almost 70% of A(H3N2) viruses circulating in North America by early 2018. Unlike most fast-growing clades, however, A2/re contains no amino acid substitutions in the hemagglutinin (HA) segment. Moreover, HI assays did not suggest substantial antigenic differences between A2/re viruses and viruses sampled during the 2016-2017 season. Rather, we observe that the A2/re clade was the result of a reassortment event that occurred in late 2016 or early 2017 and involved the combination of the HA and PB1 segments of an A2 virus with neuraminidase (NA) and other segments a virus from the clade A1b. The success of this clade shows the need for antigenic analysis that targets NA in addition to HA. Our results illustrate the potential for non-HA drivers of viral success and necessitate the need for more thorough tracking of full viral genomes to better understand the dynamics of influenza epidemics.

---

Influenza virus epidemics cause significant morbidity and mortality with about 10% of the global population infected annually [1] and most recent mortality estimates indicate that there are 291,000–645,000 influenza virus associated death each year [2]. Seasonality of influenza epidemics can vary by geographic location. In temperate regions, influenza activity typically peaks in mid-winter, while seasonal patterns are less stereotypical in tropical or subtropical regions [3]. Additionally, the severity of influenza epidemics can also vary by season and geography. The 2017–18 influenza season in the United States was the most severe epidemic since the 2003– 04 season, resulting in 79,000 deaths and more than 330,000 hospitalizations in the United States alone [4].

Throughout life humans are repeatedly infected by influenza viruses, primarily due to antigenic drift wherein amino acid substitutions in the hemagglutinin (HA) and/or neuraminidase (NA) surface glycoproteins generate viruses that escape from immunity induced by prior infection or vaccination. Out of the four co-circulating lineages of epidemic influenza viruses, A(H3N2) evolves the most rapidly, with generally higher circulation levels, necessitating more frequent updates to the specific strain used in the seasonal influenza vaccine [5, 6]. Due to ramifications in vaccine strain selection, improvements to tracking and predicting spread of A(H3N2) variants presents an opportunity for scientific insights to improve public health outcomes [7].

Research on influenza antigenic evolution focuses primarily on HA, whose antigenic properties are historically analyzed with hemagglutination inhibition (HI) assays [8]. Amino acid substitutions in epitope regions, primarily in the globular head of the HA protein, alter antigenic phenotype of the virus as measured by HI titers [5, 9–11]. However, NA evolves at nearly the same rate as HA and has similar signatures of adaptive evolution [12]. Additionally, recent work showed the importance of anti-NA antibodies to protective immunity through natural infection and vaccination [13, 14]. This suggests that NA evolution is a substantial contributor to the changing antigenic properties and dynamics of influenza viruses. While most research is devoted to the study of antigenic drift via specific amino acid substitutions, separate genomic segments coding for HA and NA can reassort, allowing novel genomic constellations to arise, a process that occurs frequently in nature [15, 16]. Reassortment can sometimes create novel adaptive genotypes [17, 18], but on average results in deleterious incompatible genotypes [19, 20]. In one notable example, the spread of the Fujian/2002 antigenic variant was attributable in part to reassortment between HA and other genomic segments, including NA [21]. The rapid development of sequencing technology and specific strategies for influenza virus genomics in the past decade enables sequencing of thousands of full viral genomes annually, where each genome consists of the coding sequences on each of the 8 influenza genome segments minus segment termini. Now nearly all influenza-positive surveillance specimens received by CDC are sequenced directly using a next generation sequencing (NGS) strategy and submitted to the GISAID Epiflu database [22]. Sequencing direct clinical samples avoids issues of virus evolution during in vitro propagation. From this abundance of genomic data, the dynamics of reassortment can be analyzed at high temporal and genotypic resolution.

Here, we use tools from the Nextstrain project [23] to show that a reassortant A(H3N2) genome constellation dominated the 20172018 North American influenza season. We go on to show that viruses with this genome constellation had the HA and PB1 segments of one parental virus with the other six segments of another virus, and that the progenitor virus likely emerged in late 2016 or early 2017.

## METHODS

### Sequence data

We built phylogenetic trees and frequency estimates for each segment of the influenza virus genome containing data from a two-year window between April 2016–April 2018, as well as a small set of older reference viruses. A total of 18,844 A(H3N2) sequences were downloaded from GISAID EpiFlu database with submission dates ranging between 2012-04-15 and 2018-04-20 (see Supplemental Text). From these, sequences were subsampled in order to achieve equal distribution of samples across geographic regions and over a two-year window (see Supplemental Text TSV for full list of samples used in analysis). In total, 1,862 sequences were used to construct phylogenetic trees.

### Phylogenetic analysis

For each segment of the influenza virus genome, alignments were created using MAFFT [24]. Aligned sequences were then cleaned of insertions relative to the reference in order to maintain consistent numbering of sites, and viruses that deviate too much from the expected molecular clock were removed. For each tree we included a set of reference viruses that are well characterized (i.e. past vaccine strains) with sampling dates before the two-year window of interest. We then constructed phylogenetic trees using FastTree [25] and used RAxML [26] to refine trees. We inferred dates and ancestral sequences of each internal node of the trees [27]. Finally, we annotated trees with attributes (i.e. geographic location), and inferred these traits for ancestral nodes.

### Reassortment analysis

To detect reassortment, we constructed tangle treessets of two trees representing different segments, with tips from matching viruses connected by lines. We then compared topologies of tangle trees that matched the phylogeny of HA segment with that of other segments. When incongruent tree topologies were found, we compared whether the viruses labelled as A2 and A2/re in the HA phylogeny were adjacent in the other segments phylogeny, and if it appeared that A2/re was a subclade of A2. If not, we inferred that a reassortment had taken place that combined the HA segment of an A2-like background with the other segment from a separate background.

## RESULTS

### Severe 2017-2018 North American influenza season

In many aspects the 2017-2018 North American influenza season was the most severe influenza epidemic in the past 12 seasons, resulting in a large increase in visits for out-patient influenza-like illness (ILI) (Fig. 1) and resulted in an estimated 960,000 hospitalizations and 79,000 influenza-associated deaths, particularly among those 65 and older [28]. In the United States there were over 23,000 confirmed cases of influenza A viruses between October 2017 and February 2018; of these 89.9% were A(H3N2) viruses. At the seasons peak, influenza A viruses were responsible for 7.7% of all hospitalizations and 10.1% of deaths in the United States [29]. The fraction of health care visits for ILI surpassed those of the 2009 influenza epidemic season (Fig. 1)—a season marked by the emergence of the A(H1N1)pdm09 pandemic virus, that continues to cause annual epidemics in humans [30]. In addition to rising to high levels, the ILI score stayed above 4% for 10 weeks during the 2017-2018 North American influenza season.

**Figure 1.**
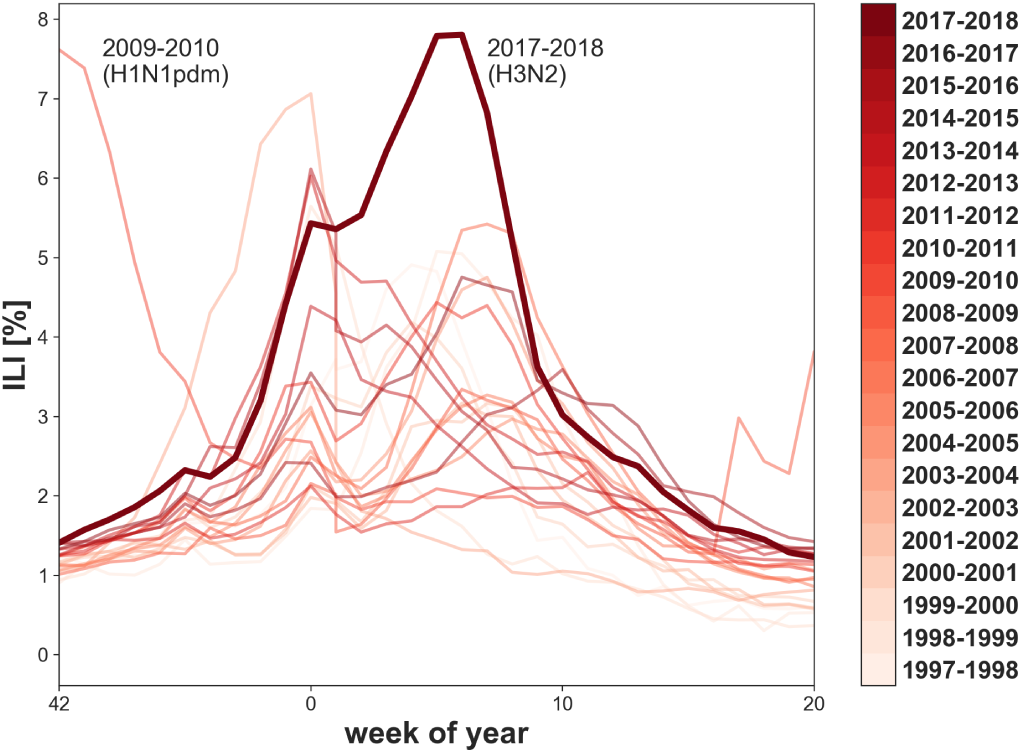
Percentage of health care visits for influenza-like illness (ILI) in the US from the 1997-1998 to the 2017-2018 season with season indicated by color. The 2017-2018 season is highlighted as a thick line; it showed the highest ILI rate since the 2009 H1N1 pandemic virus. Data from CDC (2018).

### Genetic diversity of recent A(H3N2) viruses

Here, we analyze viral clades, representing clonal descendants of a single common ancestor, on the HA segment of A(H3N2). Rapid clade turnover results in an A(H3N2) HA phylogeny in which contemporaneous viruses typically coalescence to a single common ancestor within 3 years [31]. However, the current most recent common ancestor (MRCA) of the global A(H3N2) virus population, existed more than 6 years ago in early 2012 (Fig. 2). Correspondingly, there is significant genetic diversity among the HA segments of circulating viruses, with the competing clades 3c3.A and 3c2.A comprising the deepest split in the phylogeny. Descendants of the 3c3.A viruses circulated at low frequencies in the global population between 2016 and 2018. Several HA subclades of 3c2.A, comprising 3c2.A1 to 3c2.A4, circulated at high frequencies between 2016 and 2018. Throughout the following, we drop the 3c2 to abbreviate these names to A1a, A1b, A2, A3 and A4. By April 2018, viruses in clades A1b and A2 predominated the global population, comprising an estimated 86% of A(H3N2) viruses (Fig. 2).

**Figure 2.**
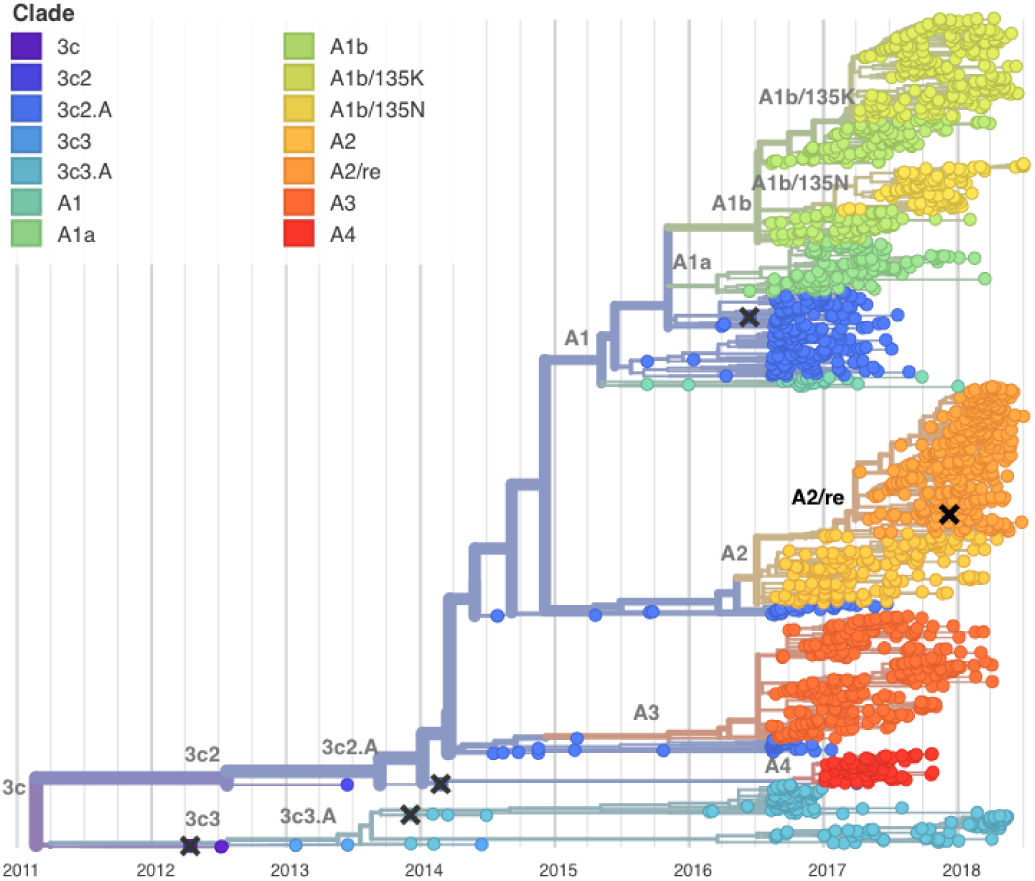
Temporally resolved phylogenetic tree of recent A(H3N2) HA sequences. Shown are influenza viruses sampled between April 2016 and April 2018 colored by clade designation (in addition, the tree shows a small number of earlier reference viruses). Viruses are subsampled to give comparable counts across time and geography. HA segments of the majority of co-circulating A(H3N2) viruses in of the 2017-2018 Northern Hemisphere influenza season fell within clades A1b (top, yellow) or A2 (center, orange). Vaccine viruses are indicated by cross marks.

Clade A1 emerged in 2015 and represented the majority of influenza A(H3N2) viruses in the 2016-2017 season. Most viruses in this clade differ from the previous vaccine virus (A/Hong Kong/4801/2014) by amino acid substitutions in HA (N121K, N171K, I406V, and G484E). Clade A1 further diversified into two subclades now named A1a and A1b. These are distinguished by the amino acid substitutions HA:G479E and HA:K92R,H311Q, respectively. Further rapid evolution of took place within A1b which split into two subclades defined by the HA amino acid substitutions E62G,R142G,T135K, and T135N. The recurrent changes at position 135 (including a loss of a putative glycosylation) suggest an adaptive origin of this rapid evolution. In the past year A1b has out-competed A1a.

Clade A2 is defined by a series of amino acid substitutions in HA:T131K, R142K, R261Q. This series of substitutions completed in mid-2016; no further amino acid changes were observed and the frequency of this clade remained fairly constant from late-2016 to mid-2017 (Fig. 3). In the 2017-2018 season in North America, however, an A2 HA subclade dominated the viruses circulating, yet lacked additional amino acid substitutions. This subcladewhich we denote as A2/re (Fig. 2) appears to have arisen as a result of a reassortment event, as discussed below. The rapid rise of subclade A2/re— coupled with an extraordinarily high incidence of that subclade in North America—stands out. By the beginning of 2018, we find the clade viruses belonging to clade A2 increasing to make up almost 70% of A(H3N2) circulating viruses, overtaking A1b as the dominant HA clade (Fig. 3).

**Figure 3.**
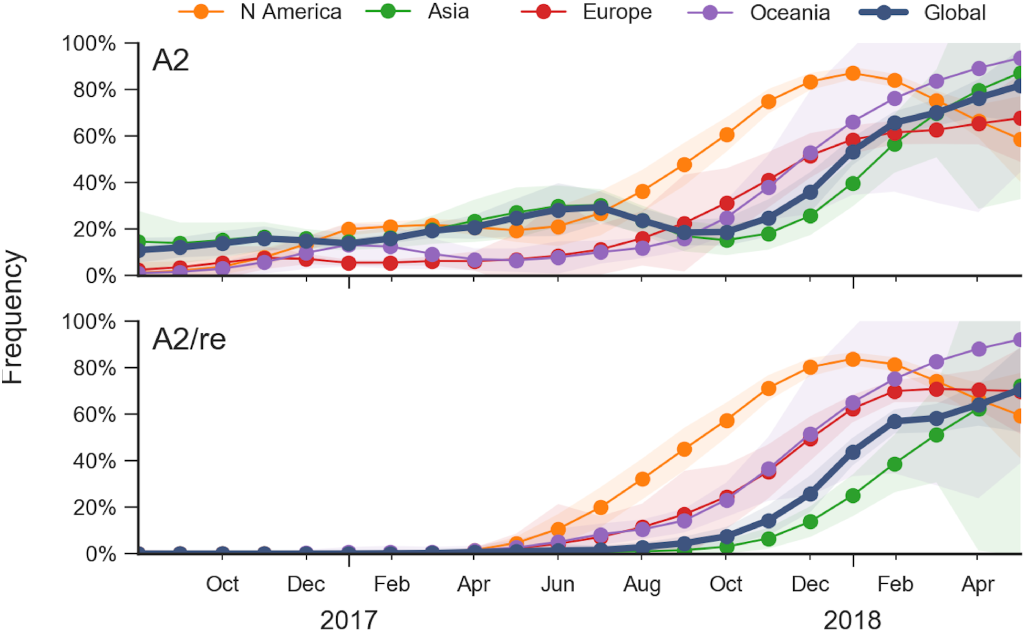
Frequency trajectories of A2 and A2/re clades partitioned by region. We estimate frequencies of different clades based on sample counts and collection dates. These estimates are based on all available data and global frequencies are weighted according to regional population size and relative sampling intensity. We use a Brownian motion process prior to smooth frequencies from month-to-month [32]. Transparent bands show an estimate the 95% confidence interval based on sample counts.

### Antigenic analysis illustrates little antigenic drift

The impact that the amino acid substitutions of the various co-circulating HA clades have on antibody escape were analyzed using post infection ferret antisera against cell-propagated viruses including vaccine strains (Supplemental Table SI). The recommended A(H3N2) vaccine viruses northern hemisphere were A/Hong Kong/4801/2014-like (i.e., A/Michigan/15/2014) for the 2016-2017 season and A/Singapore/INFIMH-16-0019/2016-like (A/Singapore/INFIMH-16-0019/2016 or A/North Carolina/4/2016) for the 2017-2018 season. Focus reduction assay of reference viruses and contemporary test viruses circulating worldwide demonstrate that antisera to either HA clade 3c2.A A/Michigan/15/2014 virus or HA clade 3c2.A1 viruses A/Singapore/INFIMH-16-0019/2016 efficiently neutralize viruses expressing 3c2.A, A1 or A2 HA RNAs (Supplemental Table SI, Supplemental Figure S1). In fact, only viruses expressing 3c3.A HAs, which represent a small proportion of viruses circulating, showed evidence of significant antigenic drift (i.e., 8-fold or greater reductions in titer as compared to titer of the homologous virus and antisera) (Supplemental Table 1). Nevertheless, antisera to the A2 HA containing virus (A/Nebraska/02/2017) efficiently neutralizes viruses containing A2 HAs and poorly neutralizes viruses expressing other HA clades (Supplemental Table 1, Supplemental Fig 1). Collectively, antigenic analysis data indicates that while the A2 HA proteins are antigenically distinct, they have not significantly drifted from 3c2.A HA proteins and it does not explain their predominance in North America during the 2017-2018 season. Consistently, substitutions to epitope sites show that the A2 clade is slightly drifted relative to ancestral 3c2.A viruses, but show little variation within clade A2 (Supplementary Fig. S1).

### Genome reassortment events among circulating viruses leads to A2/re genotype

When reassortment occurs between viruses from established clades in the HA and NA RNA segments, incongruent phylogenetic relationships are created so that a reassortment event within clade A2 viruses is immediately visible (Fig. 4). In the HA phylogeny, viruses from A2 and A2/re are closely related and form a monophyletic clade. However in the NA phylogeny, the NA of A2 viruses appear in a clade distant from the NA of A2/re viruses. Viruses whose HA belongs to clade A2/re possess NA with the N329S substitution which is shared with viruses that belong to clade A1b. The NA segments of viruses belonging to HA clade A2/re differ from viruses belonging to clade A2 by NA substitutions I176M, K220N, N329S, D339N, and P386S. Importantly, the amino acid substitution N329S is present in the NA of viruses from both successful HA clades (A2/re and A1b), as indicated by the local branch index (LBI) which summarizes recent clade growth (Supplemental Fig. S2) [33].

**Figure 4.**
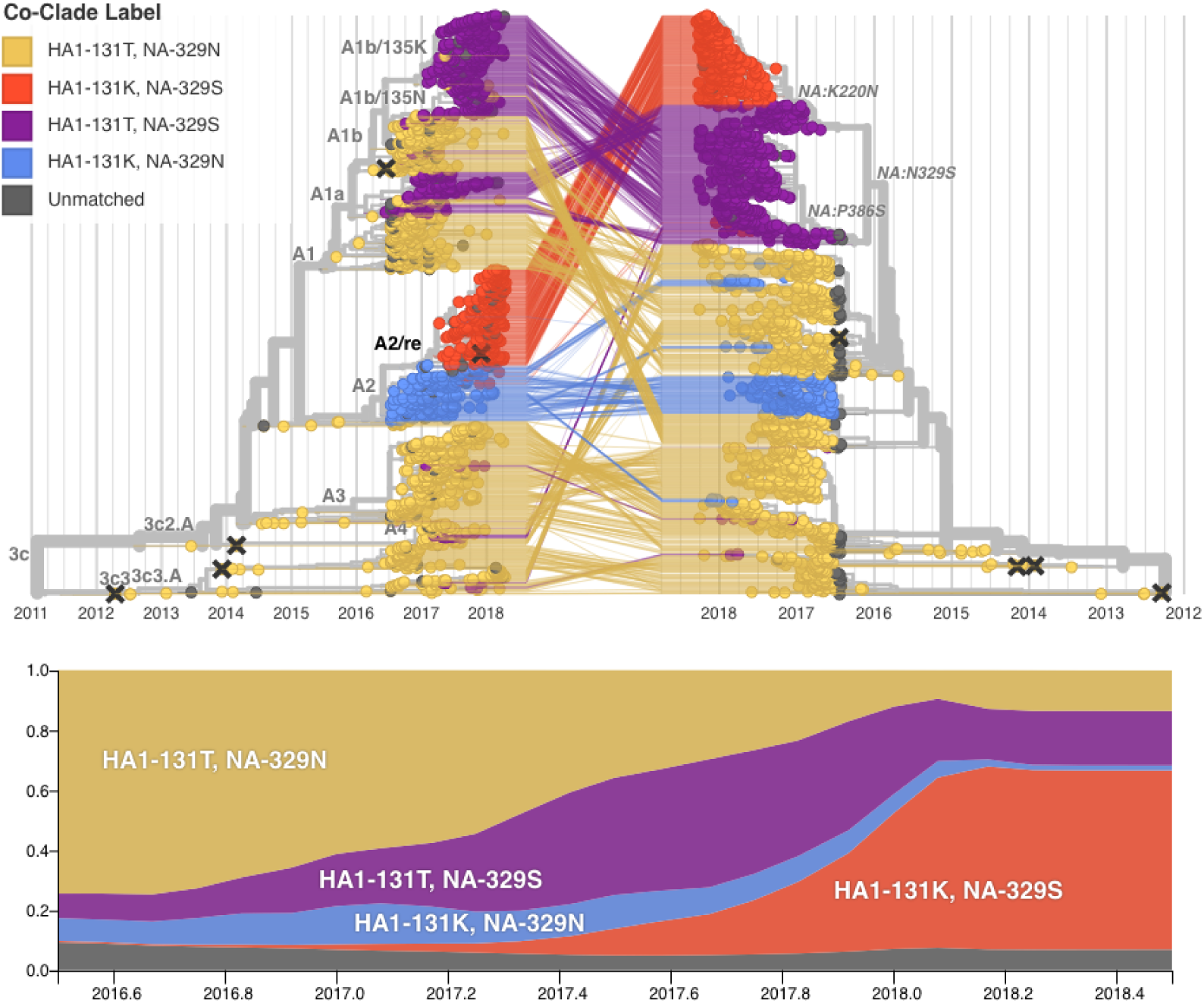
*(top)* Reassortment combines existing genetic diversity and results in incongruent phylogenetic trees. The figure shows time-scaled phylogenetic trees of HA (left) and NA (right) colored genotype at HA site 131 and NA site 329. The HA and NA sequences from the virus isolates are connected by lines. We see that the closest relatives for the viruses belonging to A2/re genotype (red) in HA are A2 viruses (blue), while their closest relatives in NA are A1 viruses (purple). *(bottom)* Global estimated frequency plots of virus clades over time. There is moderate growth of A1 viruses prior to the reassortment, but after the reassortment takes place we observe rapid growth of the reassortant clade to approximately 70% frequency. Branches on the HA tree are labelled by HA clade designation; branches on the NA tree are labelled by NA amino acid substitution. Vaccine strains are indicated by cross marks.

Temporal phylogenetic analysis places the MRCA of the A2/re genotype between Jan 2017–Feb 2017 (HA) and between Dec 2016–Feb 2017 (NA), suggesting that a reassortment event in late 2016 or early 2017 gave rise to a virus that seeded the rapidly growing population with this genome constellation. We observe that viruses basal to the reassortant A2/re clade in both HA and NA are predominantly from North and South America with A/Minnesota/12/2017, collected in February 2017, representing an early example. A discrete trait phylogeographic model places the root of the A2/re reassortment event as 83% North America / 17% South America (HA) and 100% North America (NA).

To quantify the rise of A2/re viruses, we calculated frequencies of viruses with genotypes HA:131T / NA:329N (ancestral HA, ancestral NA), HA:131T / NA:329S (ancestral HA, derived NA), HA:131K / NA:329N (derived HA, ancestral NA) and HA:131K / NA:329S (derived HA, derived NA-A2/re genotype) (Fig. 4). The three former genotypes were observed at frequencies around 10-20% in 2016, but the A2/re genotype first appeared in February 2017. By December 2017, this genotype had risen to above 70% frequency in North America; this is a comparable rate to that at which the 3c3 clade rose to predominance during 20122013.

The HA phylogeny for A2/re viruses are also incongruent compared to A2 viruses within trees for the six other RNA segments (PB2, PA, NP, NA, MP, NS). As with the NA, the six other RNA segments of A2/re viruses are most closely related to those of HA clade A1b containing viruses (Supplemental Fig. S3). Phylogenetic analysis of the PB1 shows that viruses from the reassortant clade A2/re are adjacent to A2 viruses (Supplemental Fig. S3). Upon examining all other segments for incongruent phylogenies, we find evidence that the A2/re genotype is the result of reassortment events pairing HA and PB1 segments from A2 viruses with all other segments from A1b viruses (Fig. 5).

**Figure 5.**
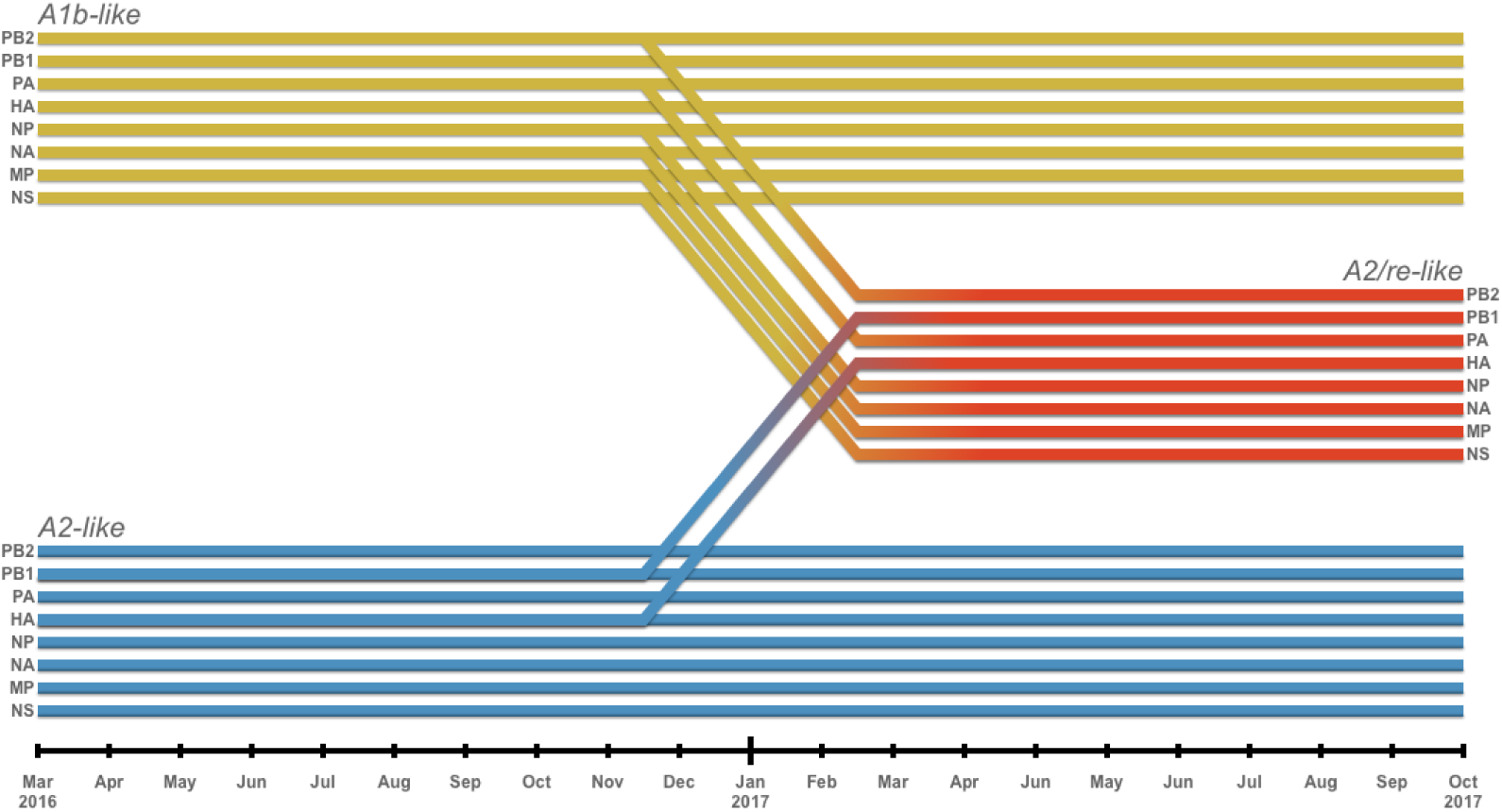
Segments from two backgrounds of A(H3N2) reassorted to form the A2/re clade (red). Segments HA and PB1 from an A2-like background (blue)combined with the six other segments from an A1b-like virus(yellow). Diagonal lines between November 2016 and February 2017 indicate the inferred timing of the reassortment.

## DISCUSSION

Most A(H3N2) influenza viruses that circulated during the 2017-2018 season in North America had hemagglutinin consensus sequences that were very similar to those of to A2 viruses, which also circulated during the previous 2016-2017 season. Consistently, antigenic analysis did not suggest significant antigenic evolution among these viruses. Only genomic sequencing and analysis of all viral RNA segments revealed that recent A2 viruses differ substantially in six of the eight RNA segments. This new genotype which we call A2/re is likely the result of a single reassortment event that took place in late 2016 or early 2017. This reassortant genotype combines the HA and PB1 segments of A2-like viruses with all other segments from an A1b-like virus. Viruses with the A2/re genotype quickly rose to high frequency within North America, coincident with one of the most severe influenza virus epidemics in recent years. During the 2018 influenza vaccine selection for the southern hemispheres 2019 season A/Switzerland/8060/2017, an A2/re virus, was selected as the A(H3N2) component of the vaccine [34].

While HA antigenic properties are well characterized across A(H3N2) viruses, antigenic characteristics of NA are not as well understood. The NA protein of these viruses differs from the previous A2 NA viruses at several positions including position 329. Position 329 changed repeatedly over the past few years and substitutions at this site could result in a loss of N-linked glycosylation. The N329S clade rose rapidly in prevalence prior to the reassortment event that gave rise to A2/re, supporting the hypothesis that the genomic background including the NA segment acquired in the reassortment event aided the success of A2/re viruses. Additionally, the NA acquired two substitutions—I176M and P386S—prior to the reassortment event. Further data are required to understand the full effect of these substitutions on NA antigenicity, both within the A2/re viruses and other genotypes. In addition to antigenic effects, replication capacity of the virus depends on interactions between different segments and their stochiometry, as has been demonstrated for different HA and NA variants [35].

While it is clear that there was rapid expansion/dissemination of A2/re genotype viruses, we cannot at this time show evidence that this clades success was solely the result of the reassortment event. Reassortment is relatively common in influenza virus evolution (as evidenced by the frequent crossing of lines in Fig. 4). Hence assigning significance to any specific reassortment event is difficult and the rapid growth of the A2/re genotype might also be driven by other epidemiological factors.

The capacity of influenza viruses to spread likely depends on the specific genome constellation of all eight RNA segments. Hence it is important to systematically survey viral evolution and reassortments of influenza viruses by genomic sequencing and a comprehensive analysis of these data. Retrospective analysis of historical data should be conducted to quantify the degree to which reassortment events give rise to successful genotypes [15, 18, 20]. Results from such analysis can then inform predictive modeling efforts to anticipate composition of future influenza virus populations [7, 33, 36, 37]. In addition to paramount importance for public health, the wealth of genomic longitudinal data with high spatiotemporal resolution make influenza viruses an ideal system to address general questions of evolutionary biology on epistasis, reassortment, and recombination.

The inclusion of genomic predictors that draw information from all eight RNA segments into fitness models will yield more robust prediction methodologies that account for more than just HA variance. We believe that integrating these sources of information to models of influenza evolution will help to predict future influenza genotype growth and us to mitigate the effect of future influenza outbreaks that would otherwise pose great risk to global human health.

## Supporting information

Metadata for sequence data.

Accession list and references for sequence data.

## Supplemental Tables and Figures

**Table SI.** Focus reduction assay of reference viruses and contemporary test viruses.

| Reference Virus | Clade | Reference Ferret Antisera |  |  |  |
| --- | --- | --- | --- | --- | --- |
|  |  | A/Michigan<br>/15/2014 | A/Singapore<br>/INFIMH-16-0019/2016 | A/Nebraska<br>/2/2017 | A/California<br>/179/2016 |
|  |  | **3c2.A** | **A1** | **A2** | **A3** |
| A/Michigan/15/2014 | 3c2.A | **640** | 320 | 1600 | 320 |
| A/Singapore/Infimh-16-0019/16 | A1 | 640 | **1280** | 800 | 1280 |
| A/Nebraska/02/2017 | A2 | 640 | 1280 | **12800** | 1280 |
| A/California/179/2016 | A3 | 640 | 2560 | 800 | **1280** |
| **Test Virus** |  |  |  |  |  |
| A/Haiti/5464/2017 | A1b | 640 | 1280 | 1600 | 1280 |
| A/Maryland/13/2018 | A1b | 320 | 1280 | 400 | 640 |
| A/Kuwait/922/2018 | A1b | 320 | 1280 | 400 | 640 |
| A/Kazakhstan/079/2018 | A2 | 640 | 1280 | 12800 | 640 |
| A/Kazakhstan/031/2018 | A2 | 640 | 640 | 12800 | 1280 |
| A/Alberta/Rv2371/2017 | A2 | 320 | 640 | 6400 | 640 |
| A/Tennessee/70/2017 | A2/re | 640 | 2560 | 25600 | 1280 |
| A/Puerto Rico/09/2018 | A2/re | 640 | 2560 | 25600 | 1280 |
| A/Peru/3817/2017 | A2/re | 640 | 640 | 12800 | 640 |
| A/Cambodia/949/2017 | A3 | 320 | 640 | 800 | 640 |
| A/New Jersey/23/2018 | 3c3.A | 80 | 320 | <200 | 80 |
| A/Delaware/20/2018 | 3c3.A | 20 | 160 | <200 | 40 |
| A/Massachusetts/18/2018 | 3c3.A | 40 | 80 | <200 | 40 |

**Figure S1.**
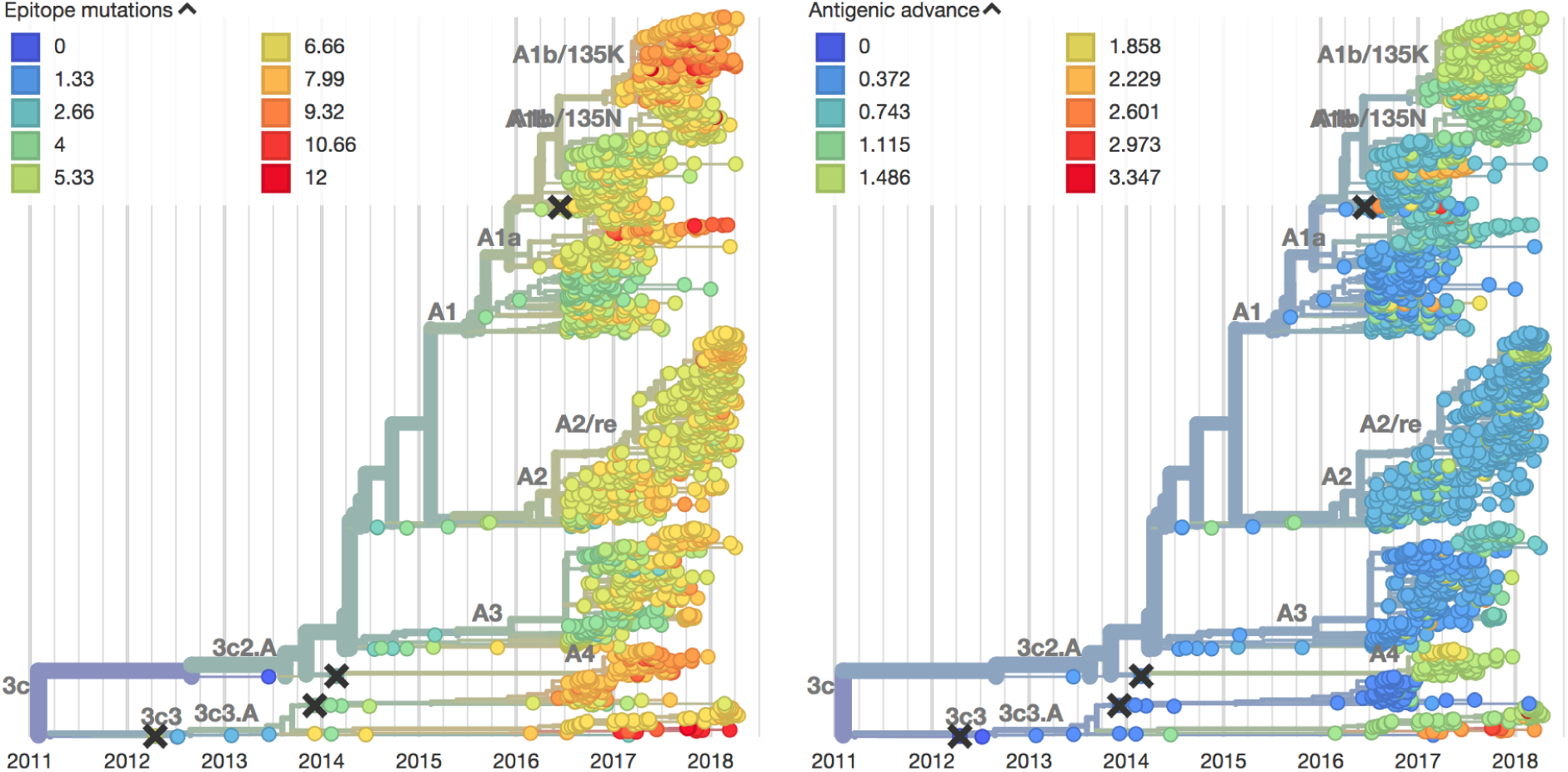
*(left)* Temporally resolved haemagglutinin phylogenetic tree colored by amino acid mutations in epitope sites [38] of HA. *(right)* The same tree, colored by antigenic advance—a measure of how much antigenic drift viruses have undergone based on related viruses’ HI titers. Both measures normally indicate that a clade may be prone to grow rapidly, due to increased ability to escape immune pressure. In both trees A2/re does not demonstrate any change from A2, making its sudden growth unusual.

**Figure S2.**
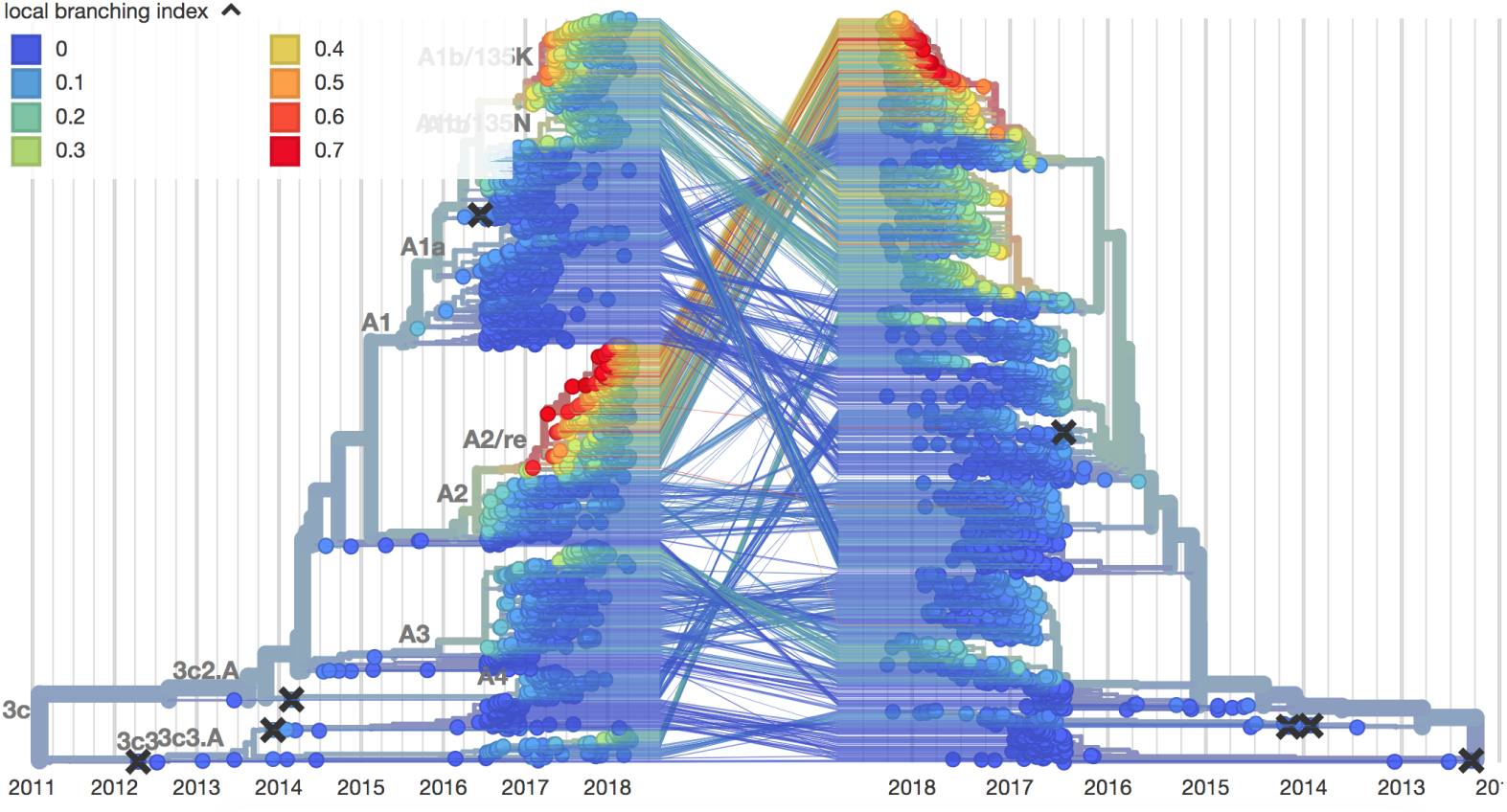
Tangle tree matching phylogenetic trees of HA and NA, colored by local branching index (LBI)—a measure that captures rapid expansion of clades. Here, LBI reveals that there is an inflection point where rapid clade growth begins at the same time that A2/re differentiates from A2 in the HA tree. Similarly, LBI is highest in the matched viruses of the NA tree, indicating that the reassortant virus saw much greater success than either of the background viruses from which it arose.

**Figure S3.**
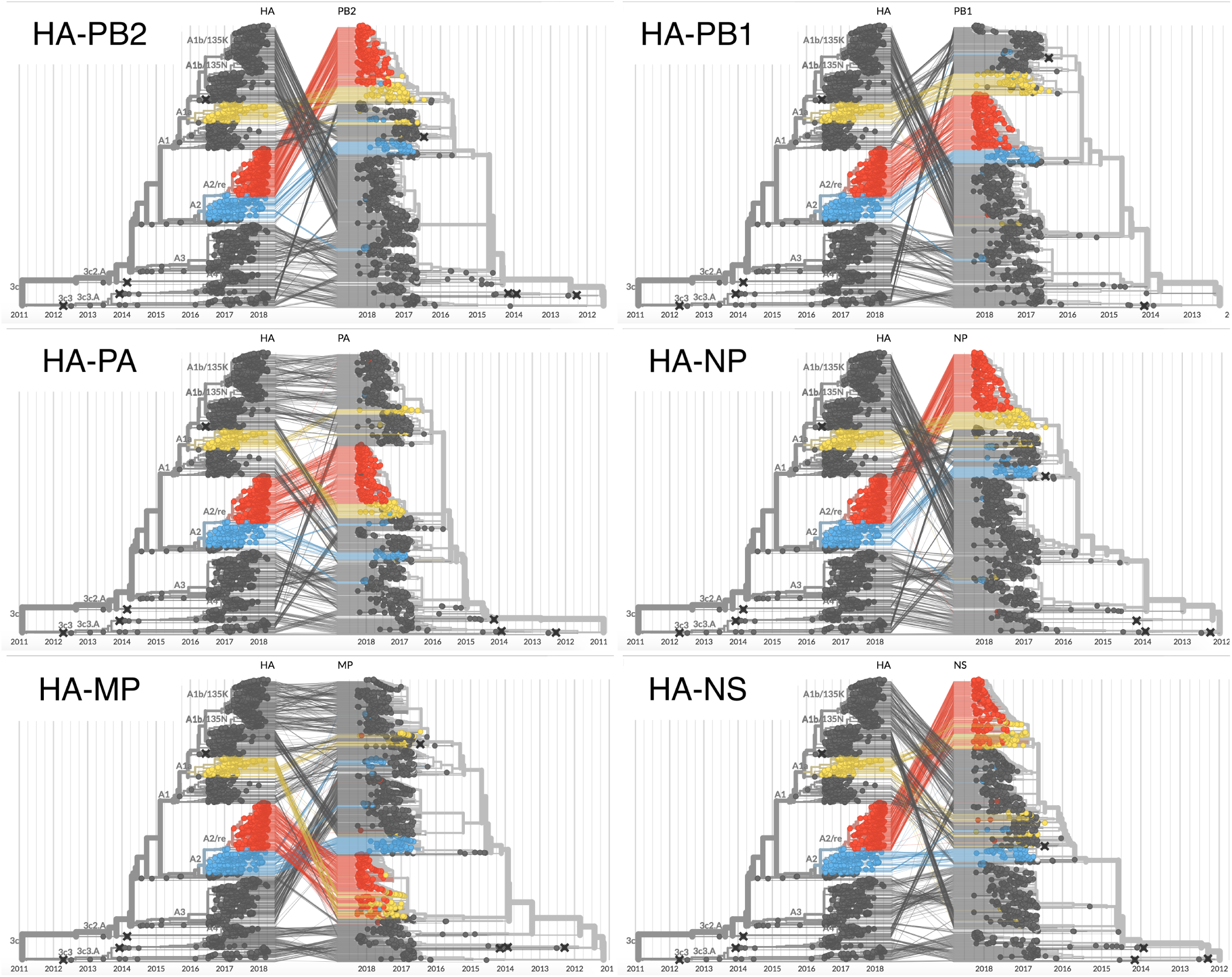
Tangle trees matching the HA phylogeny (always on left) with the six other non-NA segments. Clade A2 is colored in blue, A1b is colored in yellow, and A2/re is in red. In all segments other than PB1 (top right) A2 and A2/re lie in different parts of the tree in the second segment than in HA, indicating that reassortment has taken place.

## REFERENCES

[1] World Health Organization, “Influenza (Seasonal),” (2018).

[2] A. D. Iuliano, K. M. Roguski, H. H. Chang, D. J. Muscatello, R. Palekar, S. Tempia, C. Cohen, J. M. Gran, D. Schanzer, B. J. Cowling, et al., The Lancet 391, 1285 (2018).

[3] V. N. Petrova and C. A. Russell, Nature Reviews Microbiology 16, 47 (2018).

[4] R. Garten, L. Blanton, A. I. A. Elal, N. Alabi, J. Barnes, M. Biggerstaff, L. Brammer, A. P. Budd, E. Burns, C. N. Cummings, et al., Morbidity and Mortality Weekly Report 67, 634 (2018).

[5] T. Bedford, M. A. Suchard, P. Lemey, G. Dudas, V. Gregory, A. J. Hay, J. W. McCauley, C. A. Russell, D. J. Smith, and A. Rambaut, eLife Sciences 3, e01914 (2014).

[6] A. J. Hay, V. Gregory, A. R. Douglas, and Y. P. Lin, Philosophical Transactions of the Royal Society of London. Series B 356, 1861 (2001).

[7] D. H. Morris, K. M. Gostic, S. Pompei, T. Bedford, M. Łuksza, R. A. Neher, B. T. Grenfell, M. Lässig, and J. W. McCauley, Trends in Microbiol (2017).

[8] G. K. Hirst, J Exp Med 75, 49 (1942).

[9] D. J. Smith, A. S. Lapedes, J. C. d. Jong, T. M. Bestebroer, G. F. Rimmelzwaan, A. D. M. E. Osterhaus, and R. A. M. Fouchier, Science 305, 371 (2004).

[10] B. F. Koel, D. F. Burke, T. M. Bestebroer, S. v. d. Vliet, G. C. M. Zondag, G. Vervaet, E. Skepner, N. S. Lewis, M. I. J. Spronken, C. A. Russell, M. Y. Eropkin, A. C. Hurt, I. G. Barr, J. C. d. Jong, G. F. Rimmelzwaan, A. D. M. E. Osterhaus, R. A. M. Fouchier, and D. J. Smith, Science 342, 976 (2013).

[11] R. A. Neher, T. Bedford, R. S. Daniels, C. A. Russell, and B. I. Shraiman, Proceedings of the National Academy of Sciences of the United States of America 113, E1701 (2016).

[12] S. Bhatt, E. C. Holmes, and O. G. Pybus, Molecular Biology and Evolution 28, 2443 (2011).

[13] A. S. Monto, J. G. Petrie, R. T. Cross, E. Johnson, M. Liu, W. Zhong, M. Levine, J. M. Katz, and S. E. Ohmit, J Infect Dis 212, 1191 (2015).

[14] Q. S. Huang, D. Bandaranayake, T. Wood, E. C. Newbern, R. Seeds, J. Ralston, B. Waite, A. Bissielo, N. Prasad, A. Todd, L. Jelley, W. Gunn, A. McNicholas, T. Metz, S. Lawrence, E. Collis, A. Retter, S.-S. Wong, R. Webby, J. Bocacao, J. Haubrock, G. Mackereth, N. Turner, B. McArdle, J. Cameron, G. Reynolds, M. G. Baker, C. C. Grant, C. McArthur, S. Roberts, A. Trenholme, C. Wong, S. Taylor, P. Thomas, J. Duque, D. Gross, M. G. Thompson, M.-A. Widdowson, and SHIVERS investigation team, The Journal of Infectious Diseases (2018), 10.1093/infdis/jiy443.

[15] M. I. Nelson, C. Viboud, L. Simonsen, R. T. Bennett, S. B. Griesemer, K. S. George, J. Taylor, D. J. Spiro, N. A. Sengamalay, E. Ghedin, J. K. Taubenberger, and E. C. Holmes, PLOS Pathogens 4, e1000012 (2008).

[16] N. Marshall, L. Priyamvada, Z. Ende, J. Steel, and A. C. Lowen, PLOS Pathogens 9, e1003421 (2013).

[17] A. D. Neverov, K. V. Lezhnina, A. S. Kondrashov, and G. A. Bazykin, PLOS Genetics 10, e1004037 (2014).

[18] G. Dudas, T. Bedford, S. Lycett, and A. Rambaut, Molecular Biology and Evolution 32, 162 (2015).

[19] R. Rabadan, A. J. Levine, and M. Krasnitz, Influenza and Other Respiratory Viruses 2, 9 (2008).

[20] M. Villa and M. Lässig, PLOS Pathogens 13, e1006685 (2017).

[21] E. C. Holmes, E. Ghedin, N. Miller, J. Taylor, Y. Bao, K. St George, B. T. Grenfell, S. L. Salzberg, C. M. Fraser, D. J. Lipman, et al., PLoS Biol 3, e300 (2005).

[22] S. Elbe and G. Buckland-Merrett, Global Challenges 1, 33 (2017).

[23] J. Hadfield, C. Megill, S. M. Bell, J. Huddleston, B. Potter, C. Callender, P. Sagulenko, T. Bedford, and R. A. Neher, Bioinformatics (2018), 10.1093/bioinformatics/bty407.

[24] K. Katoh and D. M. Standley, Mol Biol Evol 30, 772 (2013).

[25] M. N. Price, P. S. Dehal, and A. P. Arkin, Mol Biol Evol 26, 1641 (2009).

[26] A. Stamatakis, Bioinformatics 30, 1312 (2014).

[27] P. Sagulenko, V. Puller, and R. A. Neher, Virus Evol 4, vex042 (2018).

[28] U. CDC, “Estimated influenza illnesses, medical visits, hospitalizations, and deaths in the united states 20172018 influenza season — cdc,” (2018).

[29] A. P. Budd, D. E. Wentworth, L. Blanton, A. I. A. Elal, N. Alabi, J. Barnes, L. Brammer, E. Burns, C. N. Cummings, T. Davis, B. Flannery, A. M. Fry, S. Garg, R. Garten, L. Gubareva, Y. Jang, K. Kniss, N. Kramer1, S. Lindstrom, D. Mustaquim, A. Ohalloran, S. J. Olsen, W. Sessions, C. Taylor, X. Xu, V. G. Dugan, J. Katz, and D. Jernigan, MMWR Morb Mortal Wkly Rep 2018, 169 (2018).

[30] R. J. Garten, C. T. Davis, C. A. Russell, B. Shu, S. Lindstrom, A. Balish, W. M. Sessions, X. Xu, E. Skepner, V. Deyde, M. Okomo-Adhiambo, L. Gubareva, J. Barnes, C. B. Smith, S. L. Emery, M. J. Hillman, P. Rivailler, J. Smagala, M. d. Graaf, D. F. Burke, R. A. M. Fouchier, C. Pappas, C. M. Alpuche-Aranda, H. Lpez-Gatell, H. Olivera, I. Lpez, C. A. Myers, D. Faix, P. J. Blair, C. Yu, K. M. Keene, P. D. Dotson, D. Boxrud, A. R. Sambol, S. H. Abid, K. S. George, T. Bannerman, A. L. Moore, D. J. Stringer, P. Blevins, G. J. Demmler-Harrison, M. Ginsberg, P. Kriner, S. Waterman, S. Smole, H. F. Guevara, E. A. Belongia, P. A. Clark, S. T. Beatrice, R. Donis, J. Katz, L. Finelli, C. B. Bridges, M. Shaw, D. B. Jernigan, T. M. Uyeki, D. J. Smith, A. I. Klimov, and N. J. Cox, Science http://dx.doi.org/10.1126/science.1176225 (2009), x10.1126/science.1176225.

[31] T. Bedford, S. Riley, I. G. Barr, S. Broor, M. Chadha, N. J. Cox, R. S. Daniels, C. P. Gunasekaran, A. C. Hurt, A. Kelso, A. Klimov, N. S. Lewis, X. Li, J. W. McCauley, T. Odagiri, V. Potdar, A. Rambaut, Y. Shu, E. Skepner, D. J. Smith, M. Suchard, M. Tashiro, D. Wang, X. Xu, P. Lemey, and C. A. Russell, Nature 523, 217 (2015).

[32] R. A. Neher and T. Bedford, Bioinformatics, btv381 (2015).

[33] R. A. Neher, C. A. Russell, and B. I. Shraiman, eLife Sciences 3, e03568 (2014).

[34] I. G. Barr, J. McCauley, N. Cox, R. Daniels, O. G. Engelhardt, K. Fukuda, G. Grohmann, A. Hay, A. Kelso, A. Klimov, T. Odagiri, D. Smith, C. Russell, M. Tashiro, R. Webby, J. Wood, Z. Ye, and W. Zhang, Vaccine 28, 1156 (2010).

[35] H.-L. Yen, C.-H. Liang, C.-Y. Wu, H. L. Forrest, A. Ferguson, K.-T. Choy, J. Jones, D. D.-Y. Wong, P. P.-H. Cheung, C.-H. Hsu, O. T. Li, K. M. Yuen, R. W. Y. Chan, L. L. M. Poon, M. C. W. Chan, J. M. Nicholls, S. Krauss, C.-H. Wong, Y. Guan, R. G. Webster, R. J. Webby, and M. Peiris, Proceedings of the National Academy of Sciences 108, 14264 (2011), http://www.pnas.org/content/108/34/14264.full.pdf.

[36] M. Łuksza and M. Lässig, Nature 507, 57 (2014).

[37] T. R. Klingen, S. Reimering, C. A. Guzmn, and A. C. McHardy, Trends in Microbiology 26, 119 (2018).

[38] Y. I. Wolf, C. Viboud, E. C. Holmes, E. V. Koonin, and D. J. Lipman, Biol. Direct 1, 34 (2006).

